# PRMT5-Mediated Arginine Methylation of HSP90AA1 Drives Esophageal Squamous Cell Carcinoma Progression

**DOI:** 10.64898/2026.03.09.710340

**Authors:** Tangbing Chen, Yilv Lv, Jing Wang, Yonggang Yuan, Kuan Xu, Minjun Shi, Wang Li, Bo Ye

## Abstract

Dysregulated HSP90AA1 chaperone activity is a hallmark of multiple human cancers, yet its post-translational modifications in esophageal squamous cell carcinoma (ESCC) are poorly defined. Here, we identify a PRMT5-driven modification of HSP90AA1, symmetric dimethylation at arginine 182 (R182), as a pivotal switch that fuels ESCC progression. HSP90AA1 physically associates with PRMT5, and genetic or pharmacologic PRMT5 blockade diminishes HSP90AA1-R182 methylation and downstream oncogenic signaling. Functionally, HSP90AA1 loss suppresses ESCC cell proliferation, migration, and invasion; enforces cell cycle arrest and reverses epithelial-mesenchymal transition (EMT). Re-expression of wild-type HSP90AA1 rescues cancer cell growth and EMT *in vitro* and *in vivo*, whereas the methylation-deficient R182A mutant does not. Furthermore, clinically oriented analysis shows HSP90AA1 overexpression in ESCC tissues and indicates moderate prognostic utility. Moreover, dual targeting of the PRMT5/HSP90AA1 axis is therapeutically attractive, and combined PRMT5/HSP90AA1 inhibition acts synergistically to exert anti-tumor activity, markedly reducing clonogenicity, invasion, and tumor burden in both cell-derived and patient-derived xenograft models. These findings demonstrate that PRMT5/HSP90AA1 serve as mechanistic drivers and actionable biomarkers in ESCC and provide a rationale for biomarker-guided co-inhibition of PRMT5/HSP90AA1 in translational studies.

## Background

Esophageal squamous cell carcinoma (ESCC) is a primary contributor to cancer-related mortality worldwide, particularly in parts of Asia and Africa where its incidence and death rates are highest [1, 2]. Challenges in early detection, combined with the disease’s rapid progression [3, 4], often result in unfavorable clinical outcomes [5–7]. Hence, advancing both the understanding and early diagnosis of ESCC remains crucial. The development of ESCC is driven by a multifaceted interplay among genetic, environmental, and epigenetic factors [8]. From a genetic standpoint, ESCC exhibits familial clustering, with variations in oncogenes such as TP53 and CDKN2A, as well as mutations in tumor suppressor genes, contributing to disease susceptibility [9–12]. Environmental determinants, including smoking, alcohol use, consumption of extremely hot foods, and vitamin deficiencies, further promote ESCC by inducing DNA damage and disrupting key cellular signaling pathways [13, 14]. Meanwhile, epigenetic alterations, notably DNA methylation and histone modifications, play integral roles in ESCC pathogenesis, often silencing tumor suppressor genes [15–18].

Heat shock protein 90AA1 (HSP90AA1) is a ubiquitously expressed molecular chaperone essential for the proper folding, stability, and function of numerous client proteins, including steroid hormone receptors, protein kinases, and transcription factors [19, 20]. It orchestrates various biological processes, ranging from stress responses and immune regulation to cell cycle control and tumor progression, highlighting its importance in maintaining cellular homeostasis under both physiological and pathological conditions [19–21]. HSP90AA1 harbors numerous residues subject to diverse post-translational modifications (PTMs), including phosphorylation, acetylation, SUMOylation, nitrosylation, O-linked N-acetylglucosamine (O-GlcNAc) modification, and ubiquitination [22]. Aberrant regulation of these modifications can instigate disease pathogenesis. For instance, dysregulated DNA methylation of the HSP90AA1 gene is associated with hepatocellular carcinoma, influencing tumor-related processes such as the cell cycle and signal transduction [23]. Moreover, differential methylation of the HSP90AA1 promoter region may underlie stress responses during pregnancy and function as a pathophysiological mechanism in preeclampsia [24]. These findings suggest that aberrant methylation of HSP90AA1 could enhance tumor cell proliferation, invasiveness, and resistance to apoptosis. Nevertheless, current investigations primarily focus on promoter-level methylation and non-methylated PTMs of HSP90AA1, leaving the function of HSP90AA1 post-translational arginine methylation in ESCC largely unexplored.

Against this backdrop, the present study aims to clarify the critical role of HSP90AA1 methylation in ESCC. Specifically, we seek to elucidate the molecular mechanisms and functional consequences of HSP90AA1 methylation in ESCC pathogenesis. We will perform comprehensive mechanistic analyses, including the identification of kinases and methyltransferases responsible for HSP90AA1 methylation, and investigate how this epigenetic modification influences its chaperone function. Through these efforts, our work aims to shed light on a potentially pivotal regulatory pathway in ESCC, with the goal of identifying novel molecular targets that could inform both diagnosis and personalized therapeutic strategies for this malignancy.

## Methods

### Single-cell transcriptome analysis

Single-cell RNA-seq data were obtained from the GEO database (GSE145370)[1]. Tumor tissues and matched adjacent normal tissues were profiled using the 10x Genomics Chromium platform. Raw count matrices were processed in Seurat[2]. Cells with fewer than 300 detected genes, more than 6,000 genes, or mitochondrial gene content greater than 10% were excluded. Gene expression data were log-normalized using the LogNormalize method with a scale factor of 10,000. Highly variable genes were identified, followed by data scaling and principal component analysis. Graph-based clustering was performed using the Louvain algorithm, and UMAP was used for visualization. Cell types were annotated based on established marker genes.

### Definition of HSP90AA1-associated epithelial states

Epithelial cells were extracted for downstream analyses. Cells were stratified into HSP90AA1-high and HSP90AA1-low groups according to the median expression level of HSP90AA1. The distribution of HSP90AA1-associated epithelial states was visualized on UMAP embeddings. Differences in HSP90AA1 expression between tumor and adjacent normal epithelial cells were evaluated using violin plots. Differentially expressed genes between HSP90AA1-high and HSP90AA1-low epithelial cells were identified using the Wilcoxon rank-sum test implemented in Seurat. Genes with an adjusted P value < 0.05 were considered significant. Gene Ontology (GO) biological process(BP) enrichment analysis was performed using the clusterProfiler package[3] to identify functional categories associated with HSP90AA1-associated epithelial states.

### Tissue Specimens

A total of 11 esophageal squamous cell carcinoma (ESCC) tissues, together with their corresponding adjacent non-tumor counterparts, were obtained from patients who underwent surgical resection at Shanghai Chest Hospital during the period from June 2022 to December 2024. Histopathological and immunohistochemical evaluations were carried out within the same institution to confirm tissue identity. Written informed consent was provided by all participants, and the entire study was conducted under approval of the institutional ethics committee. Clinical characteristics of these patients are summarized in **Table S1**. For subsequent analyses, the expression level of HSP90AA1 in tumor specimens was quantified, and the cohort was stratified into high- and low-expression groups using the median value as the cutoff.

### Cell Lines Culture

Human ESCC cell lines (ECA109, KYSE150) together with an immortalized normal esophageal epithelial line (Het-1A) were obtained from the Cell Bank of the Chinese Academy of Sciences (Shanghai, China). Cells were maintained under standard culture conditions in RPMI-1640 medium supplemented with 10% fetal bovine serum at 37 °C in a humidified incubator containing 5% CO₂. Routine monitoring for mycoplasma contamination was performed using the MycoAlert kit (Lonza), and all lines were confirmed to be negative.

### RNA Extraction and Quantitative PCR

Total RNA from both cultured cells and clinical tissue specimens was isolated using TRIzol reagent (Invitrogen, Thermo Fisher Scientific). Reverse transcription was carried out with a commercial cDNA synthesis kit (Beyotime Biotechnology, Haimen, China), following the manufacturer’s recommendations. Quantitative real-time PCR was subsequently performed on a StepOne detection system (Applied Biosystems) with SYBR Green chemistry (Roche, Mannheim, Germany). Relative expression of candidate genes was determined using the comparative 2^−ΔΔ^ CT approach, with GAPDH serving as the internal control for normalization.

### Western Blotting

Cells were seeded into 6-well plates at a density of 1 × 10⁵ cells/mL (2 mL/well) and allowed to attach for 24 h. Following incubation, the culture medium was removed, and the cells were rinsed with PBS. Cellular proteins were extracted by lysing cells on ice for 15 min in RIPA buffer supplemented with PMSF and phosphatase inhibitors. Lysates were subsequently sonicated (1 min) and centrifuged at 12,000 rpm for 15 min at 4 °C, and the clarified supernatant was stored at −80 °C. Protein concentration was measured using a BCA assay. Equal amounts of protein were mixed with 5× SDS loading buffer, denatured at 95 °C for 5 min, and preserved at −20 °C until use.

For electrophoresis, both stacking and separating SDS-PAGE gels were freshly prepared and allowed to polymerize (10–30 min). Approximately 10 μL of protein lysate together with 3 μL of marker were loaded per lane. Electrophoresis was performed at 80 V for the stacking phase (20–30 min), followed by 120 V until the dye front reached the gel bottom. Proteins were electro-transferred to PVDF membranes at 200 mA for 1–3 h. After transfer, membranes were blocked with 5% non-fat milk in TBST for 2 h at room temperature, washed, and incubated overnight at 4 °C with primary antibodies. After additional washes, membranes were exposed to secondary antibodies for 90 min at room temperature, then visualized using ECL reagents. Signals were captured with an automated chemiluminescence imaging system.

### Histology and immunohistochemical staining

Paraffin-embedded sections were first dewaxed in xylene (three changes, 10 min each), followed by sequential rehydration through graded ethanol solutions (100%, 95%, and 75%) and final rinsing in distilled water. Slides were then immersed in PBS for 15 min. Antigen retrieval was achieved by heating in citrate buffer in a microwave oven for 5 min, repeated twice with a 20-min cooling interval. Endogenous peroxidase activity was quenched using a blocking reagent at room temperature for 10 min. After PBS washes, sections were incubated overnight at 4 °C with the primary antibody (1:50 dilution). Following additional washes, membranes were treated with an HRP-conjugated goat anti-mouse/rabbit IgG secondary antibody at 37 °C for 15 min. Visualization was carried out using freshly prepared DAB substrate for 5 min, and nuclei were counterstained with hematoxylin (20 s) followed by alcohol differentiation. For analysis, digital images of stained slides were captured with the Super Image morphological system. Three randomly selected fields at 40× magnification were recorded for each case. Quantitative assessment of positive (yellow-brown) staining was performed using ImageJ software, and results were expressed as the percentage of stained area (%Area).

### Immunofluorescence staining

Cells in logarithmic growth phase from different treatment groups were seeded into confocal culture dishes at a density ensuring 70–80% confluence. After complete adherence, culture medium was removed and cells were gently rinsed with PBS. Fixation was performed with 4% paraformaldehyde for 30 min, followed by three PBS washes (10 min each). Permeabilization was achieved with 0.5% Triton X-100 for 5 min, and sections were again rinsed with PBS. Non-specific binding was blocked by incubation with 5% goat serum for 1 h at room temperature. Cells were then incubated overnight at 4 °C with primary antibodies diluted to the manufacturer’s recommended concentration. After three washes with PBS, fluorophore-conjugated secondary antibodies were applied for 2 h in the dark at room temperature. Nuclei were counterstained with DAPI for 3 min, and after final PBS rinses, samples were imaged under a fluorescence microscope.

### Xenograft Model

To construct in vivo experiment, 5×10^6^ cells were resuspended Matrigel. The supernatant of the cell suspension corresponding to 5.5×10^6^ cells was carefully aspirated, and then the cells were gently resuspended on ice with 1.1 mL of Matrigel, avoiding generating excessive bubbles that would affect the next injection step. For subcutaneous injection, the skin of the head and neck of the nude mouse was grasped with the left thumb, and the hind legs and tail of the nude mouse were fixed with the left index finger and ring finger to fully expose the skin on the right back. The pre-injection area was disinfected with an alcohol swab. The syringe was held with the right hand, and the needle was inserted into the skin at a 45° angle downward, and then advanced parallel under the skin in the direction of the head for 5 - 10 mm. Appropriate back-and-forth movement of 3 - 5 mm was performed under the skin to ensure that there was no connective tissue at the injection site that would affect the injection effect while avoiding the needle from puncturing the subcutaneous tissue and entering the abdominal cavity. 100 μL of the Matrigel cell suspension was subcutaneously injected into each nude mouse. After the needle was withdrawn, a dry swab was quickly used to gently press the injection site to avoid the spillage of the cell suspension. On the 20 th day of inoculation, the body weight of the nude mice was measured. The nude mice were deeply anesthetized and sacrificed by cervical dislocation. The tumor tissues were dissected, the diameter of the tumor was evaluated, and the tumor volume was calculated. The volume calculation method was: 1/2 × (length × width^2^). The tumor tissues were paraffin-embedded and sectioned.

### Cell proliferation assay (CCK-8)

Cells were seeded into 96-well plates at a density of 5 × 10³ cells per well and allowed to adhere for 24 h in a humidified incubator (37 °C, 5% CO₂). Following transfection with either control shRNA or HSP90AA1-specific shRNA, cell viability was assessed at 6, 24, 48, and 72 h. At each time point, 10 μL of CCK-8 reagent was added to each well and incubated according to the manufacturer’s instructions. Absorbance at 450 nm was measured using a microplate reader. All experiments were independently repeated at least three times.

### Cell cycle analysis by flow cytometry

For cell-based experiments, adherent cells were gently detached using trypsin solution without EDTA to avoid potential over-digestion, which could otherwise generate false-positive signals. The harvested cells were subsequently washed twice with phosphate-buffered saline (PBS), and the concentration of each suspension was adjusted to approximately 1 × 10⁶ cells/mL. One milliliter of this suspension was again rinsed twice with ice-cold PBS before being resuspended in 1× binding buffer. From each sample, 100 μL of the prepared suspension was transferred into a 5-mL culture tube, after which 5 μL of fluorescein isothiocyanate (FITC)-labeled Annexin V and 5 μL of propidium iodide (PI) were added sequentially. The mixtures were incubated at ambient temperature in the absence of light for 30 min, followed by the addition of 400 μL of binding buffer. All manipulations were carried out carefully to avoid vigorous pipetting, thereby minimizing mechanical damage to the cells. After staining, samples were maintained on ice and subjected to flow cytometric analysis within 1 h to ensure stability of fluorescence signals. For each specimen, a minimum of 10,000 events was recorded. FITC-Annexin V fluorescence was detected in the FL1 channel, whereas PI signals were monitored in the FL2 channel. To define proper gating strategies, a negative control group containing unstained cells was included for every run.

### Transwell migration and invasion assays

Cell motility was evaluated using 24-well Transwell inserts equipped with polycarbonate membranes (8-μm pore size; Corning, USA). For invasion experiments, the upper surfaces of the membranes were coated with 50 μL of Matrigel (BD Biosciences, USA) diluted 1:8 in serum-free RPMI-1640 medium, followed by incubation at 37 °C for 4 h to allow gelling. Migration assays were conducted in chambers without Matrigel coating. ESCC cell lines (ECA109 and KYSE150) were first deprived of serum for 12 h, trypsinized, and resuspended in serum-free RPMI-1640. Approximately 5 × 10⁴ cells in 200 μL of medium were added to the upper compartments, while the lower chambers were filled with 600 μL of RPMI-1640 supplemented with 20% fetal bovine serum to serve as a chemoattractant. After incubation at 37 °C for 24 h (migration) or 48 h (invasion), cells remaining on the upper side of the filter were gently removed with a cotton swab. Cells that had traversed to the underside of the membrane were fixed in 4% paraformaldehyde for 20 min, stained with 0.1% crystal violet for 15 min, and then rinsed in PBS. Five randomly selected microscopic fields per membrane were photographed using an inverted microscope (Nikon Eclipse Ti, Japan) at 100× magnification. Cell numbers were quantified with ImageJ software (version 1.53, NIH). All assays were independently repeated three times to ensure reproducibility.

### Co-Immunoprecipitation (Co-IP)

ESCC cells were plated in 10-cm dishes until reaching suitable confluence, after which total protein was isolated using a lysis solution containing 20 mM Tris (pH 7.4), 150 mM NaCl, 2 mM EDTA, 2 mM EGTA, 1 mM sodium orthovanadate, 50 mM sodium fluoride, 1% Triton X-100, 0.1% SDS, and 100 mM PMSF. The lysates were clarified by centrifugation at 13,000 rpm for 10 min at 4 °C. Pre-clearing was performed with protein A/G beads for 1 h at 4 °C to minimize nonspecific binding. Afterward, 5 μg of specific primary antibody or isotype-matched IgG was added to the supernatants and incubated overnight at 4 °C with gentle agitation. On the following day, protein A or G agarose beads were introduced and further incubated for 3 h at 4 °C to capture immune complexes. The bead-bound materials were washed three times with buffer consisting of 100 mM NaCl, 50 mM Tris (pH 7.5), 0.1% NP-40, 3% glycerol, and 100 mM PMSF. Proteins retained on the beads were eluted in 2× Laemmli sample buffer, heated at 95 °C for 5 min, and subsequently analyzed by standard western blotting procedures.

### Prognostic Analysis of HSP90AA1 in ESCC

RNA sequencing data processed by the STAR pipeline for the TCGA-ESCA (esophageal carcinoma) cohort were retrieved from the TCGA database (https://portal.gdc.cancer.gov). Expression matrices in TPM format and corresponding clinical annotations were extracted and organized for subsequent analysis. Patients were stratified into high- and low-expression subgroups according to HSP90AA1 transcript levels. Differential expression analysis was performed on the raw count matrix using the DESeq2 package, with threshold criteria defined as |log₂FC| > 1 and p < 0.05. To evaluate prognostic significance, the R “survival” package was applied to conduct proportional hazards assumption testing and single-gene Cox regression analysis. Survival outcomes were visualized with the survminer and ggplot2 packages. In addition, time-dependent ROC curve analysis was carried out using the “timeROC” package, and corresponding graphics were generated with ggplot2 to compare 1-, 3-, and 5-year survival outcomes between groups defined by HSP90AA1 expression. Batch survival regression modeling was further implemented to identify additional ESCA-associated prognostic markers. Pairwise correlation analyses between HSP90AA1 and other transcripts across the dataset were then conducted. Overlapping and distinct correlation patterns were illustrated through ggplot2 and VennDiagram visualizations.

### Immune Microenvironment Analysis

To obtain a comprehensive overview of the immune contexture in ESCC, both immune cell infiltration and immune checkpoint gene expression were systematically evaluated. For estimation of immune cell subsets, two complementary deconvolution strategies were applied: CIBERSORT and QUANTISEC. The CIBERSORT algorithm, which is based on support vector regression, was employed to infer the relative abundance of 22 immune cell populations using the LM22 reference matrix. Prior to analysis, RNA-seq or microarray expression data were normalized with either TPM (Transcripts Per Million) or RMA (Robust Multi-array Average) and uploaded to the CIBERSORT portal (https://cibersort.stanford.edu/). To confirm the robustness of these results, the QUANTISEC R package was additionally used, which estimates immune infiltration on the basis of predefined cell-type gene signatures. Proportions of immune cells generated from both tools were visualized through bar plots to facilitate cross-sample comparisons. For the evaluation of immune checkpoint activity, transcript levels of major checkpoint regulators such as PD-L1, CTLA-4, and PD-1 were extracted from RNA-seq or microarray datasets. Normalization was again performed using TPM or RMA methods, and statistical tests were applied to assess expression differences with corresponding p-values. The expression profiles were displayed in bar plots and heatmaps, enabling visualization of immune checkpoint heterogeneity and offering insights into potential mechanisms of tumor immune evasion in ESCC.

### GO and KEGG Functional Enrichment Analysis

GO analysis was conducted through the clusterProfilerGO.R package within R software (https://www.r-project.org/) and the Perl language to identify common targets. Concurrently, KEGG Pathway enrichment analysis was executed using the clusterProfilerKEGG.R package. The Pathview package was utilized to draw the corresponding signaling pathway diagrams. The degree of core pathway enrichment was analyzed in line with the enrichment factor value, and the potential biological functions and signaling pathway mechanisms of core genes were probed.

### Gene Regulatory Analysis

Based on the core genes identified through the afore-mentioned procedures, the websites of TRANSFAC and JASPAR databases are accessed to predict the transcription factors on the marker genes. JASPAR (http://jaspar.genereg.net/): JASPAR is a commonly utilized database of transcription factor binding sites, providing a considerable number of sequences of transcription factor binding sites and related information on transcription factors. TRANSFAC (https://bioinfo.uth.edu/TRANSFAC/): TRANSFAC is a database encompassing transcription factor binding sites and regulatory modules of transcription factors, offering an extensive amount of experimentally verified information on transcription factor binding sites. PROMO is an online tool for predicting transcription factor binding sites, capable of making predictions based on the sequences provided by users and the transcription factor database. TRED (http://rulai.cshl.edu/TRED/): TRED is a transcription factor regulatory database that provides regulatory relationships between transcription factors and target genes and predicts transcription factor binding sites by analyzing the upstream sequences of target genes. These websites can facilitate the prediction of the transcription factors upstream of target genes and offer information and databases related to transcription factors and binding sites. Eventually, the common transcription factors predicted from the above four databases are integrated as the subsequent validation targets. Additionally, the binding site information between transcription factors and genes is obtained from the GTRD website (http://gtrd.biouml.org/). GTRD (Gene Transcription Regulation Database) was a database specifically collecting and organizing the binding site data between transcription factors and genes. In GTRD, the binding site information can be retrieved by inputting the predicted names of transcription factors and genes.

### Patient-derived xenograft (PDX) model

All procedures involving animals were carried out under protocols reviewed and approved by the Institutional Animal Care and Use Committee of Shanghai Chest Hospital, in accordance with national guidelines for laboratory animal welfare. Freshly resected tumor fragments from ESCC patients were immediately immersed in ice-cold serum-free RPMI-1640 medium (Gibco, Cat# 11875-093) and maintained at 4 °C. To preserve viability, implantation into mice was performed within 48 h after collection. For xenotransplantation, immunodeficient NPI mice were obtained from Beijing IDMO Co., Ltd. All animals were maintained under specific-pathogen-free (SPF) conditions in individually ventilated cages. The housing environment was strictly controlled with constant temperature (22 ± 1 °C), relative humidity (50 ± 1%), and a 12-h light/dark cycle. Standard rodent chow and autoclaved water were provided ad libitum. Animals were monitored daily for health status, body weight, and tumor growth, ensuring compliance with humane endpoints established by the institutional guidelines.

### Statistical Analysis

All statistical evaluations were conducted using GraphPad Prism software (version 5.0). Quantitative data are presented as mean values accompanied by standard deviation (SD). For comparisons between categorical or continuous variables, chi-square tests were applied where appropriate, while differences between two independent groups were assessed with a paired Student’s t-test. In the case of more than two groups, one-way analysis of variance (ANOVA) was carried out, followed by Tukey’s multiple-comparison post hoc test to determine pairwise significance. A p-value less than 0.05 was regarded as statistically significant in all analyses.

## Results

### Elevated HSP90AA1 Expression and Prognosis in ESCC Databases

Single-gene analysis showed significant overexpression of HSP90AA1 in esophageal cancer (p < 0.05; **Supplementary Figure 1A**). Although higher HSP90AA1 levels were associated with poorer survival, the correlation was not statistically significant (p > 0.05; **Supplementary Figure 1B**). Time-dependent ROC analysis showed a 5-year AUC of 0.764 for HSP90AA1 (**Supplementary Figure 1C**), indicating moderate prognostic value. Differential expression analysis identified 969 DEGs, with 289 upregulated and 680 downregulated (**Supplementary Figure 1D-E**). Additionally, 601 genes were linked to survival, and 5,448 genes co-expressed with HSP90AA1. Five genes (GAL, GLI2, PTGDS, XCR1, LEMD1) were common across these datasets (Supplementary Figure 1F). Analysis of pathological stages showed that these genes were enriched in advanced stages (**Supplementary Figure 1G**). These results align with HSP90AA1 expression and its association with poor prognosis, suggesting its role in tumor progression and metastasis. GO and KEGG enrichment revealed these genes were involved in processes like cerebellar granule cell proliferation, glucocorticoid metabolism, and peptide receptor activity (**Supplementary Figure 1H**).

Single-cell RNA sequencing analysis identified major cell types in tumor and adjacent normal tissues, including epithelial cells, fibroblasts, endothelial cells, and various immune cell populations (**Figure 1A-B**). Among these, epithelial cells exhibited significant transcriptional heterogeneity and were selected for further analysis. Notably, HSP90AA1 expression was unevenly distributed across epithelial cells, with significantly higher levels observed in tumor-derived epithelial cells compared to adjacent normal epithelial cells (**Figure 1C-D**), indicating the presence of an HSP90AA1-associated epithelial state. Epithelial cells from the tumor group were extracted and stratified into HSP90AA1-high and HSP90AA1-low groups based on median expression levels. GO biological process enrichment analysis revealed that HSP90AA1-high epithelial cells were predominantly enriched for processes related to protein folding, protein complex assembly, intracellular protein localization, RNA processing, and protein stability (**Figure 1E**). To further investigate whether the observed functional differences reflected coordinated regulatory programs, a set of genes related to post-translational modifications (PTM) was evaluated at the single-cell level. Compared to HSP90AA1-low cells, HSP90AA1-high epithelial cells exhibited significantly higher PTM program scores (**Figure 1F**), supporting the notion that protein-level regulatory processes are broadly enhanced in this epithelial state. Interestingly, classical oncogenic signaling pathways were not significantly enriched. Instead, the enriched biological processes were primarily related to post-transcriptional and post-translational regulation of protein handling, suggesting that HSP90AA1-associated epithelial heterogeneity is accompanied by extensive remodeling of protein regulatory programs. Given the enrichment of PTM-related programs, we examined the expression patterns of major protein arginine methyltransferase (PRMT) family members. While most PRMT genes showed low or heterogeneous expression across different epithelial states, PRMT5 exhibited significantly higher and more consistent expression in HSP90AA1-high epithelial cells (**Figure 1G, Supplementary Figure 1I**). In contrast, other PRMT family members, including PRMT1, PRMT3, PRMT7, and PRMT9, did not show similar cell state-specific enrichment.

**Figure 1.**
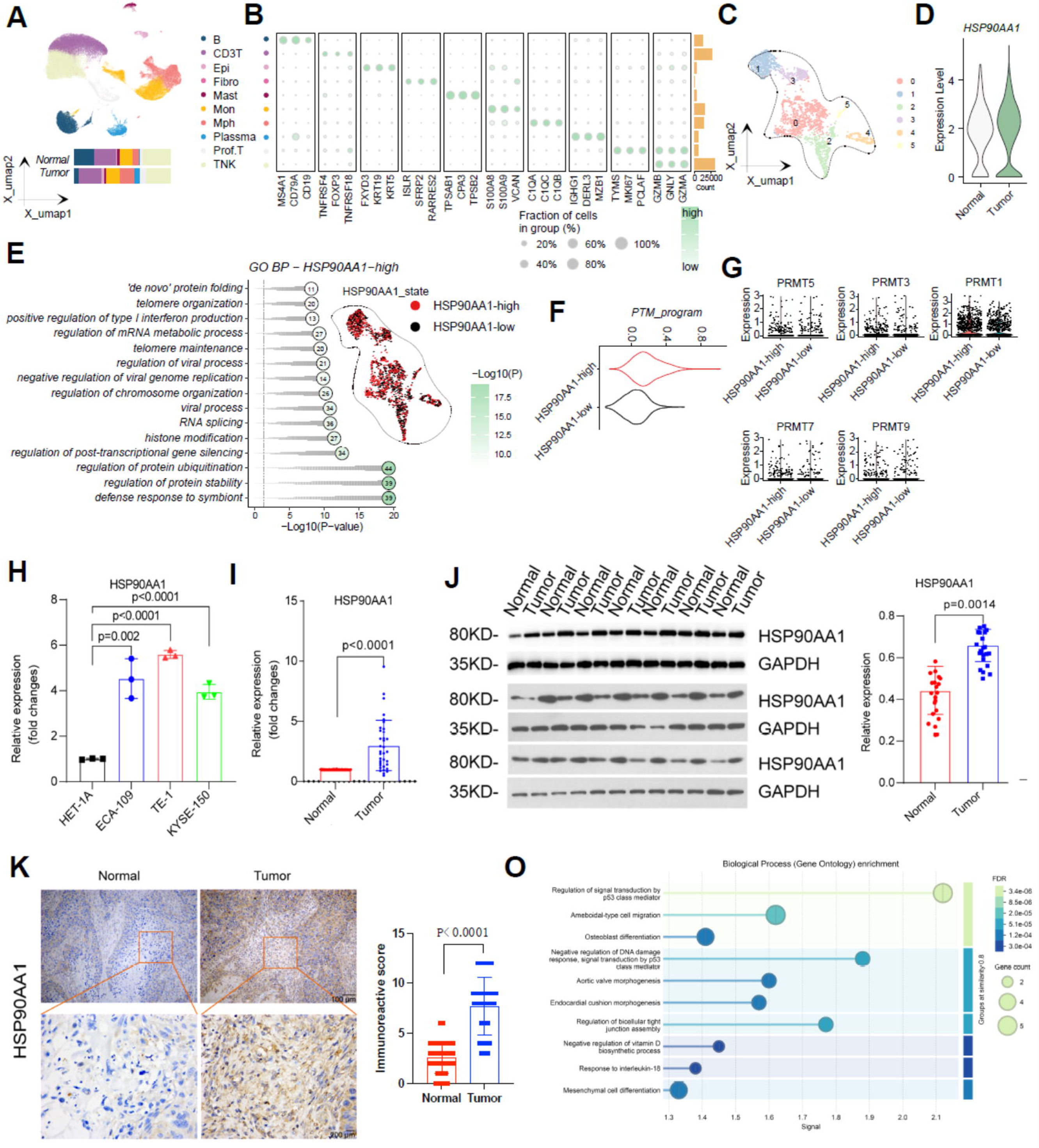
Single-cell characterization of HSP90AA1-associated epithelial states and PTM programs in ESCC: (A) UMAP visualization of large cell population clusters in ESCC, with color-coded cell types including HSP90AA1-associated populations. (B) Dot plot representing the fraction of cells expressing selected markers among different celltypes. (C) UMAP clustering of epithelial cells. (D) Violin plot comparing HSP90AA1 expression between normal and tumor tissues. (E) GO enrichment analysis of biological processes associated with HSP90AA1-high epithelial cells. (F) PTM program scores in HSP90AA1-high versus HSP90AA1-low epithelial cells. (G) Expression levels of PRMT family members (PRMT5, PRMT3, PRMT1, PRMT7, PRMT9) in HSP90AA1-high and HSP90AA1-low cells. (H) HSP90AA1 mRNA expression in normal esophageal cells (HET-1A) and ESCC cell lines (ECA-109, TE-1, KYSE-150). (I) HSP90AA1 mRNA expression in normal and tumor tissues from ESCC patients. (J) Western blot showing HSP90AA1 protein levels in normal and tumor tissues. (K) Immunohistochemical analysis confirming increased HSP90AA1 protein expression in ESCC tumor tissues compared to normal tissues. (O) GO enrichment analysis of HSP90AA1-associated proteins, highlighting key biological processes in ESCC progression.

### Immune Cell Composition and Checkpoint Molecule Expression in ESCC Tumors

To investigate HSP90AA1’s role in modulating the immune microenvironment, we used CIBERSORT and QUANTISEC to quantify immune cell composition in ESCC tumors. **Supplementary Figure 1J** (CIBERSORT) and **Supplementary Figure 1J** (QUANTISEC) show significantly increased proportions of M1 macrophages and activated NK cells, alongside reduced levels of regulatory T cells (Tregs) and B cells change suggestive of a pro-inflammatory, anti-tumor immune state. Immune checkpoint analysis (**Supplementary Figure 1K**) revealed elevated PD-L1 (CD274) expression, implicating a possible immune escape mechanism in which HSP90AA1 enhances PD-L1-mediated T-cell inhibition. Conversely, the relatively low expression of CTLA-4 and PD-1 suggests that these pathways may be less pivotal in the current context **Supplementary Figure 1L**. We next employed JASPAR, TRANSFAC, PROMO, and TRED databases to predict transcription factors for the core intersecting genes (GAL, GLI2, PTGDS, XCR1, and LEMD1). Similarity between the gene sequences and known transcription factor binding sites was used to pinpoint potential regulators. Additionally, data from the GTRD website validated these predictions. GAL, for instance, was predicted to bind SP1, while PTGDS may interact with KLF6, YY1, and NF-κB. Detailed transcriptional regulatory network was shown in **Supplementary Figure 1M**. Collectively, these findings point to a dual modulatory function for HSP90AA1: it may simultaneously foster tumor immune evasion via PD-L1 upregulation and shape anti-tumor immunity by altering immune cell composition.

### HSP90AA1 is Overexpressed in ESCC Cells and Tissues

We then confirmed HSP90AA1 overexpression at both mRNA and protein levels in ESCC cell lines and clinical tissues. **Figure 1H** showed markedly elevated HSP90AA1 mRNA in ESCC cell lines (TE-1, ECA109, and KYSE-150) compared with normal esophageal epithelial cells (HET-1A). Similarly, **Figure 1I** indicated significantly higher HSP90AA1 mRNA levels in ESCC tumor tissues than in corresponding normal tissues. Immunoblotting (**Figures 1J**) and immunohistochemistry (**Figure 1K**) further corroborated these findings, revealing a substantial increase in HSP90AA1 protein expression in ESCC tumor samples relative to normal esophageal epithelium. Furthermore, functional annotations for HSP90AA1-associated proteins indicated significant participation in regulation of signal transduction by p53 class mediator, negative regulation of DNA damage response, signal transduction by p53 class mediator, regulation of bicellular tight junction assembly and etc (**Figure 1O**).

### Knockdown of HSP90AA1 Inhibits ESCC Cell Proliferation, Metastasis, and Invasion

Knocking down HSP90AA1 using shRNA significantly decreased ESCC cell proliferation over 72 h, as determined by CCK-8 assays (**Figure 2A**). Flow cytometric cell cycle analysis (**Figures 2B** and **2C**) showed a striking reduction in S phase populations following HSP90AA1 knockdown, with a concomitant increase in G0/G1. Pharmacological inhibition of HSP90AA1 using PU-29F yielded a similar cell cycle profile (**Figures 2D** and **2E**). Western blot analysis revealed marked reductions in key cell cycle proteins (Cyclin B, PCNA) in HSP90AA1-depleted or -inhibition cells (**Figure 2F**), consistent with G0/G1 accumulation. *In vivo*, stable knockdown of HSP90AA1 or pharmacological inhibition of HSP90AA1 using PU-29F in ESCC xenograft models significantly suppressed tumor growth and final tumor weight (p < 0.001; **Figures 2G - I**), highlighting HSP90AA1’s essential role in ESCC tumorigenesis. Immunofluorescence (IF) analysis provided further mechanistic insights into HSP90AA1-mediated EMT and metastatic processes. HSP90AA1 depletion enhanced membrane-localized E-cadherin expression (**Supplementary Figure 2A**) but diminished Vimentin levels (**Supplementary Figure 2B**). Moreover, PCNA expression was markedly reduced in HSP90AA1-knockdown cells (**Supplementary Figure 2C**), supporting the antiproliferative effect observed. The EMT-driving transcription factors ZEB1 (**Supplementary Figure 2D**), β-catenin (**Supplementary Figure 2E**), and Slug (**Supplementary Figure 2F**) were significantly downregulated in HSP90AA1-knockdown cells, reflecting a transition away from a mesenchymal, pro-metastatic phenotype. Western blot analysis corroborated these findings, showing reduced N-cadherin, Vimentin, Fibronectin, Snail, Slug, ZEB1, and Twist1 upon HSP90AA1 inhibition (**Supplementary Figure 2G**). Immunohistochemical (IHC) staining and western blot analysis in tissue sections further confirmed that HSP90AA1 knockdown leads to restored epithelial markers (E-cadherin, α-catenin, γ-catenin) and decreased mesenchymal markers (Vimentin, Fibronectin, N-cadherin) (**Supplementary Figure 3**), supporting the mIF data. Together, these observations underscore HSP90AA1’s pivotal role in promoting EMT and metastasis in ESCC.

**Figure 2.**
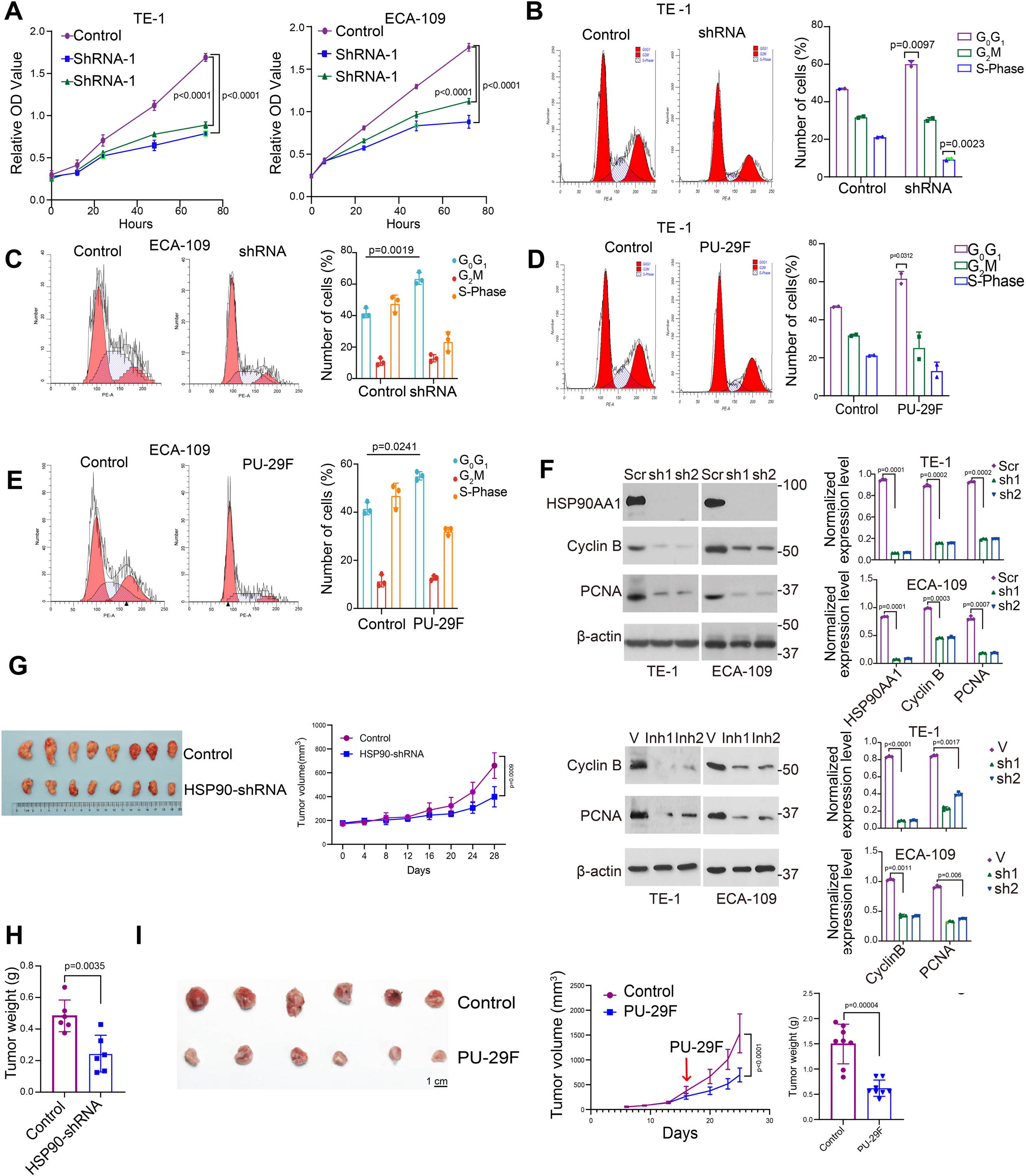
Changes in cell proliferation and cell cycle distribution after *hsp90aa1* knockdown or inhibition: (A) CCK-8 assays showing reduced proliferation in ESCC cells (ECA-109 and TE-1) with shRNA-mediated HSP90AA1 knockdown compared to controls. (B and C) Flow cytometric cell cycle analysis demonstrating a decrease in the S phase in shRNA-treated ESCC cells. (D and E) Similar cell cycle shifts (G0/G1 accumulation) were observed upon treatment with the HSP90 inhibitor PU-29F, compared to controls. (F) Western blot analysis showing reduced Cyclin B and PCNA protein levels in HSP90AA1-depleted cells, in line with G0/G1 arrest. β-actin serves as a loading control. (G and H) Impact of HSP90AA1 knockdown on tumor growth in an ESCC xenograft model. (Left) Growth curves over time; (Right) Significant decrease in final tumor weight (p < 0.001) compared to control. (I) Impact of HSP90AA1 inhibition on tumor growth in an ESCC xenograft model. (Left) Growth curves over time; (Right) Significant decrease in final tumor weight (p < 0.001) compared to control.

### PRMT5-Mediated Methylation of HSP90AA1 at Arginine 182 Enhances Its Activity in ESCC

To elucidate the molecular basis of HSP90AA1 function in ESCC, we examined its interaction with the arginine methyltransferase PRMT5 in 293T cells. Co-IP demonstrated that PRMT5 and HSP90AA1 could interact with each other (**Figure 3A**), a finding validated by parallel co-IP assays in ESCC cells (**Figure 3B**). Knocking down PRMT5 or using a PRMT5 inhibitor significantly reduced the HSP90AA1-interacted symmetric dimethylarginine (SDMA) levels in 293T cells (**Figures 3C** and **3D**), highlighting PRMT5’s major role in modulating HSP90AA1 methylation. Bioinformatic analyses identified a putative methylation site at arginine 182 (R182) of HSP90AA1 (**Figure 3E**). Site-directed mutagenesis at this locus (R182A) dramatically decreased SDMA modification in both 293T and ESCC cells (**Figures 3F** and **3G**), confirming R182 as critical for HSP90AA1’s methylation-dependent functionality. In summary, these findings reveal a novel regulatory mechanism wherein PRMT5-mediated arginine methylation of HSP90AA1 at R182 augments its activity, probably contributing to ESCC progression.

**Figure 3.**
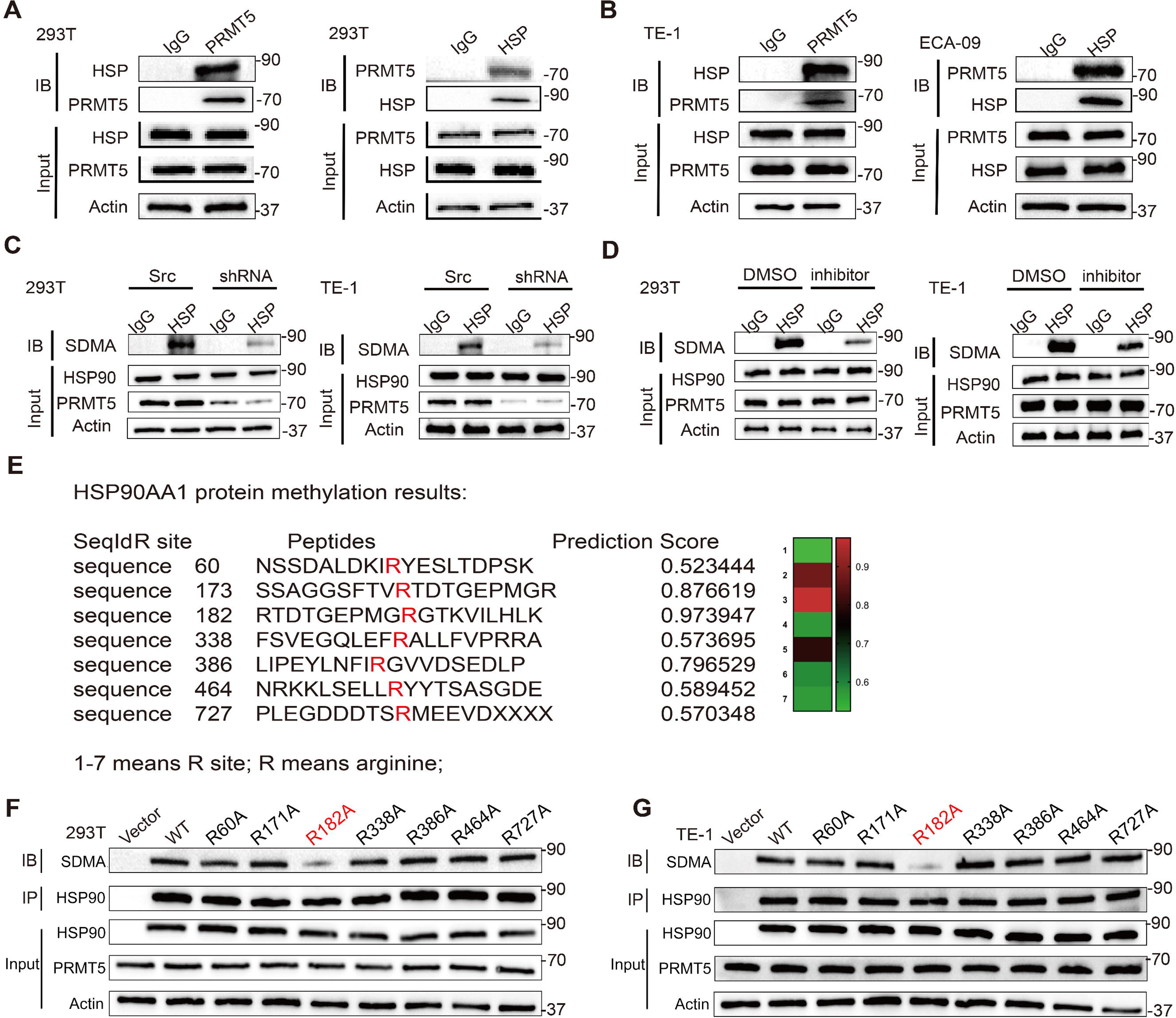
PRMT5-mediated methylation of HSP90AA1 at arginine 182 in ESCC: (A and B) Co-immunoprecipitation (Co-IP) assays in 293T, and ESCC cells demonstrated a stable interaction between HSP90AA1 and PRMT5. (C and D) Co-immunoprecipitation (Co-IP) assays in 293T indicated a reduction of the HSP90AA1-PRMT5 interaction upon HSP90AA1 knockdown or inhibition. (E) Bioinformatic identification of potential methylation sites at R60, R173, R182, R338, R386, R464, and R727 in HSP90AA1. (F and G) Co-IP analysis showing symmetric dimethylarginine (SDMA) modification in wild-type (WT) HSP90AA1 and various mutants (R60A, R171A, R182A, R338A, R386A, R464A, R721A) in 293T (F) and ESCC (G) cells, highlighting R182 as a crucial methylation site for HSP90AA1 function.

### HSP90AA1 promotes the progression of ESCC dependent on PRMT5-mediated methylation

Subsequently, we constructed HSP90AA1 knockout ESCC cells using CRISPR-Cas9 technology and confirmed its knockout efficiency (**Figure 4A**). To investigate whether the oncogenic function of HSP90AA1 relies on PRMT5-mediated methylation, we rescued HSP90AA1-depleted cells with either wild-type (WT) HSP90AA1 or the methylation-defective R182A mutant. Immunoblotting verified comparable expression levels of exogenous WT and R182A proteins (**Figure 4B**). Functional analyses revealed that reintroduction of WT HSP90AA1 fully restored cellular colony formation, and migratory capacity in knockout cells, as evidenced by colony formation assays (**Figure 4C**), scratch wound healing (**Figure 4D**), and transwell migration assays (**Figure 4E**). In contrast, cells expressing the R182A mutant exhibited significantly impaired rescue effects. Consistently, WT HSP90AA1, but not the R182A mutant, reversed the loss of mesenchymal markers (N-cadherin, Vimentin) and epithelial-marker (E-cadherin) downregulation in knockout cells (**Figure 4F**), confirming its essential role in maintaining EMT progression. *In vivo* xenograft models further corroborated these findings: tumors derived from WT-rescued cells displayed increasing growth rates and metastatic potential to control groups, whereas R182A-expressing tumors showed little effect (**Figure 4G-H**). Collectively, these data demonstrate that the R182 methylation site, a PRMT5 substrate, is indispensable for HSP90AA1’s oncogenic functions in ESCC progression.

**Figure 4.**
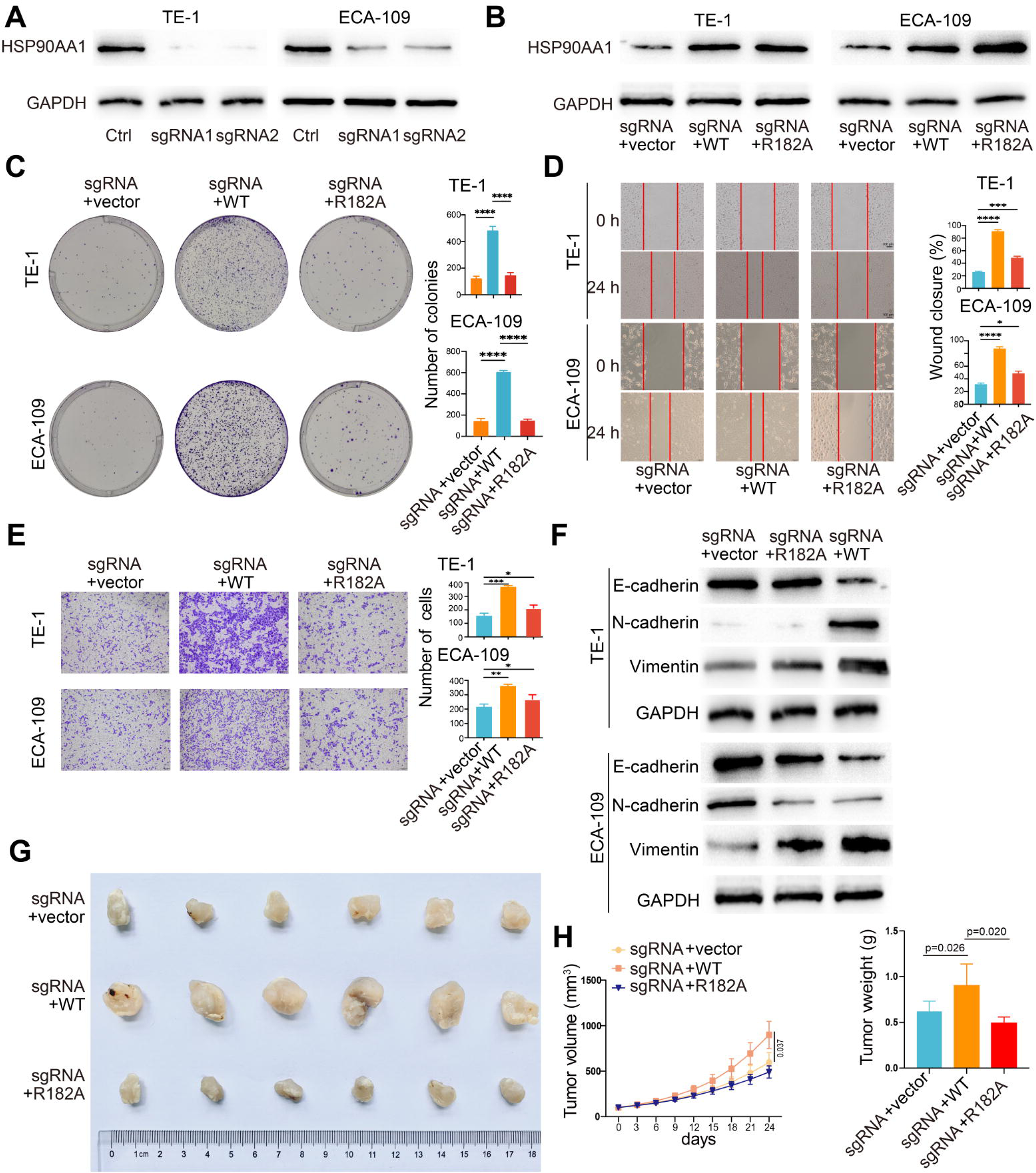
HSP90AA1 promotes the progression of ESCC dependent on PRMT5-mediated methylation. (A) The knockout efficiency of sgRNA against HSP90AA1. (B) The expression of HSP90AA1 in ESCC cells with HSP90AA1 knockout plus WT form of HSP90AA1 overexpression or HSP90AA1 R182A overexpression. (C) The ability of clone formation of ESCC cells was evaluated in ESCC cells described in (B). (D) The migration ability of ESCC cells was determined in ESCC cells depicted in (B). (E) The invasion ability of ESCC cells was evaluated in ESCC cells described in (B). (F) The expression of EMT markers was detected in ESCC cells described in (B). (G and H) Impact of WT HSP90AA1 or R182A HSP90AA1 rescuing on tumor growth in an ESCC xenograft model. (Left) Tumor images derived from cells; (Right) Growth curves over time and significant decrease in final tumor weight (p < 0.001) compared to control.

### The combinatory inhibition of PRMT5 and HSP90AA1 exhibits significant suppression on ESCC progression in PDX model

To validate the therapeutic potential of targeting the PRMT5-HSP90AA1 axis, we established patient-derived xenograft (PDX) models using freshly resected ESCC tumors implanted subcutaneously in immunodeficient mice. After 28 days of treatment, the combination group demonstrated markedly reduced tumor growth compared to monotherapy groups (**Figure 5A**), with final tumor volumes decreased by 72.3% (vs. control, *p*<0.001) versus 41.2% and 38.7% reductions in PRMT5i and HSP90i groups, respectively (**Figures 5B - 8D**). Molecular analysis of PDX tumors revealed profound EMT modulation by combination therapy. IHC analysis demonstrated robust E-cadherin restoration at cell-cell junctions and dramatic reductions in mesenchymal markers Snail and Twist (**Figure 5E**). Western blotting further confirmed coordinated upregulation of epithelial markers (E-cadherin, γ-catenin) and downregulation of EMT transcription factors (Slug, Twist1, ZEB1) and matrix components (fibronectin) in the combination group (**Figure 5F**), indicating potent suppression of EMT *in vivo*. These findings were corroborated by diminished HSP90AA1 methylation and PRMT5 activity (symmetric dimethylarginine levels) in combination-treated tumors (**Figure 5G**) confirming mechanistic synergy between PRMT5 and HSP90AA1 inhibition. Collectively, these PDX model results demonstrate that dual targeting of PRMT5 and HSP90AA1 synergistically inhibits ESCC progression by simultaneously blocking oncogenic methylation signaling and reversing EMT-driven metastatic programs.

**Figure 5.**
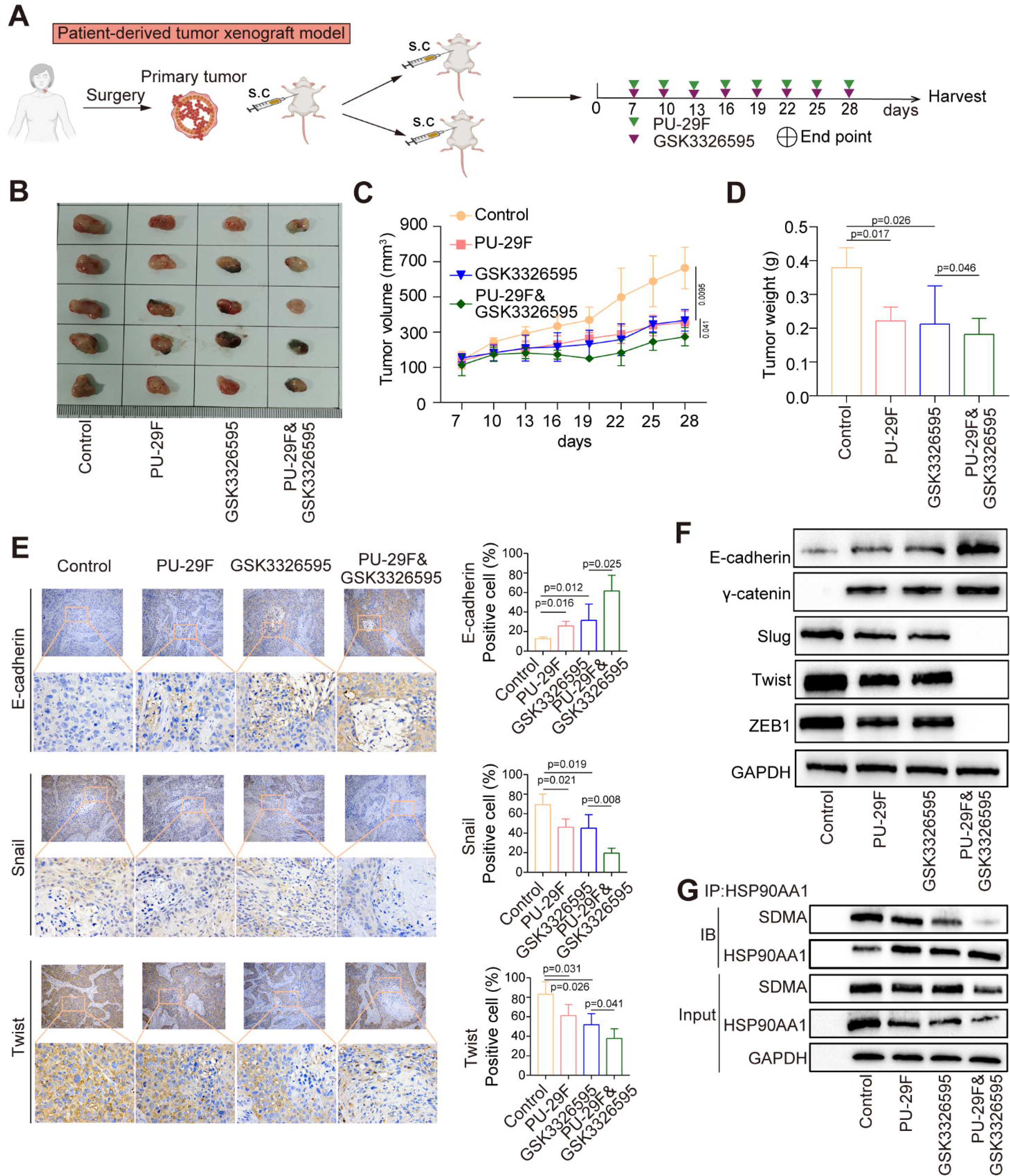
The combinatory inhibition of PRMT5 and HSP90AA1 exhibits significant suppression on ESCC progression in PDX model. (A) PDX model was established to evaluate the combinatory inhibition of PRMT5 and HSP90AA1 on ESCC progression. (B) Tumor images derived from PDX model. (C and D) Growth curves over time (C) and significant decrease in final tumor weight (D). (E) The expression of EMT markers was examined in tumor tissues derived from the PDX model using IHC assay. (F) The expression of EMT markers was detected in tumor tissues derived from the PDX model using WB analysis. (G) Co-IP analysis showing symmetric dimethylarginine (SDMA) modification of HSP90AA1 in tumor tissues from the PDX model.

### Synergistic targeting of HSP90AA1 and PRMT5 induces potent anti-tumor effects in PDX model

To further evaluate the combinatorial efficacy of HSP90AA1 and PRMT5 inhibition, ESCC cell line ECA109 was treated with escalating doses of HSP90AA1 inhibitor AUY922 and PRMT5 inhibitor GSK3326595. Cell proliferation analysis revealed strong synergistic effects in inhibitory concentrations reduced by 6.8-to 9.3-fold compared to monotherapies (**Figure 6A**, *p*<0.001). Colony formation assays demonstrated near-complete eradication of clonogenic potential under combination treatment (*p*<0.001; **Figure 6B**), while transwell assays showed reductions in invasion (*p*<0.01; **Figure 6C**). Flow cytometry revealed a striking reduction in S phase populations (*p*<0.001) in combination-treated cells (**Figure 6D**). In subcutaneous xenografts, combination therapy suppressed tumor growth by 83.6% (vs. 51.2% and 47.8% for AUY922 and GSK3326595 alone; *p*<0.001; **Figure 6E - G)**. Additionally, combination treatment synergistically reversed EMT, as evidenced by the upregulation of E-cadherin and reductions in N-cadherin, vimentin, Slug, and ZEB1 (*p*<0.001; **Figure 6H**), paralleled by diminished symmetric dimethylarginine (SDMA) modification of HSP90AA1 (R182me2s) and PRMT5 activity (**Figure 6I)** These data establish that dual inhibition of HSP90AA1 and PRMT5 exerts synergistic anti-tumor effects by concurrently blocking proliferation, metastatic dissemination, and EMT programs in ESCC.

**Figure 6.**
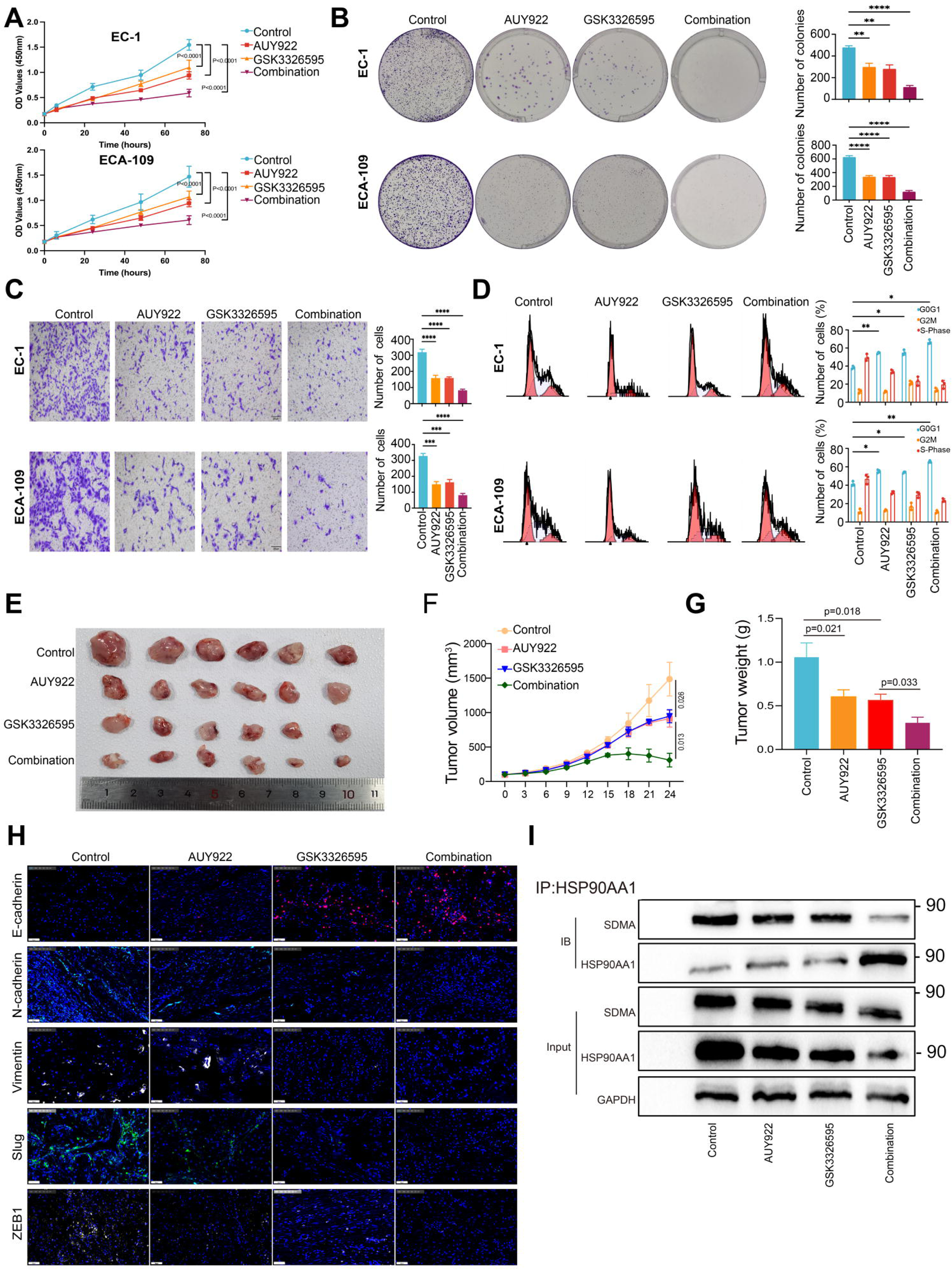
Synergistic targeting of HSP90AA1 and PRMT5 induces potent anti-tumor effects in CDX model. (A) CCK-8 assays showing reduced proliferation in ESCC cells (ECA-109 and TE-1) with inhibition of HSP90AA1 or PRMT5, individually, or combinative inhibition of them compared to controls. (B) Clone formation assays indicating reduced proliferation in ESCC cells (ECA-109 and TE-1) with inhibition of HSP90AA1 or PRMT5, individually, or combinative inhibition of them compared to controls. (C) Transwell-invasion analysis demonstrating reduced invasion in ESCC cells (ECA-109 and TE-1) with inhibition of HSP90AA1 or PRMT5, individually, or combinative inhibition of them compared to controls. (D) Flow cytometric cell cycle analysis demonstrating a decrease in the S phase ESCC cells (ECA-109 and TE-1) with inhibition of HSP90AA1 or PRMT5, individually, or combinative inhibition of them compared to controls. (E - G) Impact of inhibition of HSP90AA1 or PRMT5, individually, or combinative inhibition of them on tumor growth in an ESCC xenograft model. (E) Tumor images derived from the CDX model; (F) Growth curves over time; (G) Significant decrease in final tumor weight (p < 0.001) compared to control. (H) The expression of EMT markers was determined in tumor tissues derived from the CDX model. (I) Co-IP analysis showing symmetric dimethylarginine (SDMA) modification of HSP90AA1 in tumor tissues from the CDX model.

## Discussion

In this study, we provided the first demonstration that PRMT5-mediated arginine dimethylation of HSP90AA1 at residue R182 plays a central role in promoting ESCC progression. By uncovering this post-translational modification (PTM), our findings expand the understanding of HSP90AA1’s multifaceted regulatory mechanisms in tumor biology. Notably, the stable interaction between HSP90AA1 and PRMT5, coupled with the specific methylation at R182, enhances the invasive and metastatic potential of ESCC cells. These observations not only underscore the significance of PTMs in orchestrating HSP90AA1 functionality but also shed light on new molecular targets for potential therapeutic intervention. Taken together, PRMT5-driven HSP90 methylation promotes EMT and tumor progression in ESCC, while HSP90 and/or PRMT5 inhibition suppresses EMT, leading to tumor regression and favorable prognosis.

As illustrated in **Figure 7**, PRMT5-driven methylation of HSP90AA1 augments its oncogenic functions, a conclusion supported by our knockdown and pharmacological inhibition data, which show that disrupting HSP90AA1 or PRMT5 substantially reduces ESCC growth and metastatic spread. These insights align with earlier research highlighting HSP90AA1’s broad involvement in cancer progression, including its roles in EMT, cell cycle regulation, migration, and invasion [29–31]. For instance, HSP90AA1 has previously been implicated in modulating key oncogenic pathways such as EGFR and HER2, contributing to tumor aggressiveness [32–33]. Our study advances this knowledge by identifying a novel methylation-dependent mechanism that further enhances HSP90AA1’s tumor-promoting activities. In contrast to earlier studies focusing predominantly on HSP90AA1 overexpression, our findings emphasize the importance of specific methylation events for sustaining the malignant phenotype of ESCC. The discovery that PRMT5 catalyzes dimethylation of HSP90AA1 at R182 reveals an additional regulatory layer with significant clinical implications. The discovery that PRMT5 catalyzes dimethylation of HSP90AA1 at R182 introduces an additional regulatory layer with marked clinical significance. By pinpointing a precise PTM that propels cancer cell proliferation, EMT, and metastasis, this work enriches the existing HSP90AA1 regulatory framework and suggests novel strategies for targeted therapeutic development.

**Figure 7.**
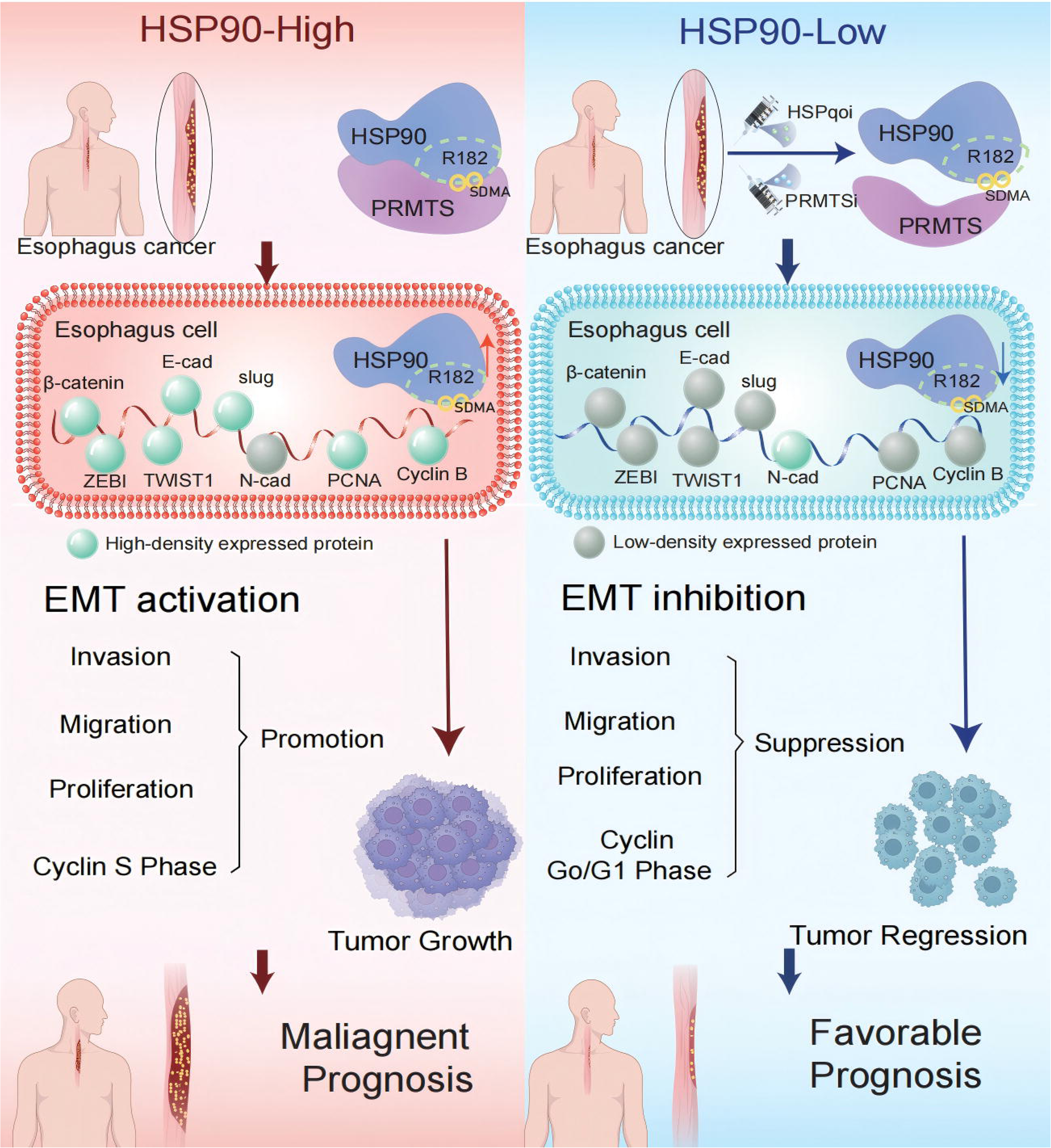
Graphic abstract. PRMT5-mediated arginine methylation of HSP90AA1 (at R182) enhances its oncogenic chaperone activity, leading to epithelial–mesenchymal transition (EMT) activation, tumor growth, and poor prognosis in esophageal squamous cell carcinoma (ESCC). In the HSP90-high state, overexpression of β-catenin, TWIST1, ZEB1, N-cadherin, PCNA, and Cyclin B promotes invasion, migration, and proliferation, driving malignant progression. Conversely, pharmacological inhibition of HSP90 (HSP90i) and/or PRMT5 (PRMT5i) diminishes HSP90 methylation, suppresses EMT, reduces tumor cell proliferation (via Cyclin Go/G1 arrest), and induces tumor regression, ultimately improving clinical outcomes.

Despite these advances, certain limitations remain. First, we conducted only a preliminary evaluation of the relationship between HSP90AA1 methylation status and clinicopathological features of ESCC and have not fully assessed its impact on patient prognosis. Future studies involving larger clinical cohorts are warranted to determine how HSP90AA1 methylation correlates with disease progression and survival outcomes, thereby strengthening its translational potential. Second, although our work focuses on PRMT5-mediated methylation, other PTMs—such as acetylation and phosphorylation—may similarly influence HSP90AA1 activity. Investigations into these modifications, as well as interactions with additional partnering proteins, will be essential for developing a more comprehensive model of HSP90AA1 regulation in ESCC pathogenesis.

Heat Shock Protein 90 (HSP90) plays a critical role in tumorigenesis when dysregulated or overexpressed, making it an attractive therapeutic target. However, its posttranslational modifications in ESCC remain poorly defined. Protein arginine methyltransferase 5 (PRMT5), responsible for di-methylating both histone and non-histone proteins, orchestrates multiple oncogenic processes including tumor growth and metastasis. Here, we demonstrate that HSP90AA1, a principal component of the HSP90 complex, physically associates with PRMT5, which catalyzes the di-methylation of HSP90AA1 at Arginine 182 (R182) in ESCC cells and tissues. This modification regulates the expression of key epithelial–mesenchymal transition (EMT) transcription factors, β-catenin, E-cadherin, Slug, PCNA, ZEB1, and TWIST1, thereby promoting ESCC cell proliferation, migration, and invasion. Notably, pharmacological inhibition of HSP90AA1 markedly suppresses tumor growth and metastasis in ESCC models. Clinically, these findings highlight the potential for co-targeting HSP90AA1 and PRMT5 as a novel therapeutic strategy, and suggest that HSP90AA1 methylation status may serve as a valuable biomarker for disease progression and treatment response. Collectively, our work uncovers a PRMT5-driven, HSP90AA1-dependent mechanism of ESCC malignancy, underscoring the importance of exploring HSP90AA1-PRMT5 inhibitors in future clinical applications.

## Conclusions

Our findings demonstrate that PRMT5-mediated methylation of HSP90AA1 at R182 drives ESCC progression by promoting EMT, cell proliferation, migration, and invasion. Further, HSP90AA1 contributes to immune evasion, in part through PD-L1 modulation, thus enhancing tumor survival. Notably, targeting the HSP90AA1–PRMT5 axis leads to significant anti-tumor effects, underscoring their therapeutic potential. These results enrich our understanding of the epigenetic regulation of HSP90AA1 and its dual role in tumor growth and immune escape, paving the way for innovative targeted therapies in ESCC. These insights deepen our understanding of the epigenetic regulation of HSP90AA1 and its dual role in tumor growth and immune escape, paving the way for innovative targeted therapies in ESCC.

## Declarations

### Authors’ contributions

T.C., Y.L., and Y.Y. contributed equally to this work and share first authorship. T.C. and Y.L. were primarily responsible for study conception, experimental design, and data collection. Y.Y. contributed to methodology optimization, data analysis, and interpretation of results. K.X. and M.S. assisted in performing experiments, data validation, and figure preparation. W.L. provided critical clinical expertise, supervised data acquisition, and contributed to manuscript revision. B.Y. supervised the overall project, provided conceptual guidance, and finalized the manuscript. All authors read and approved the final version of the manuscript.

### Funding

This work was supported by the Interdisciplinary Program of Shanghai Jiao Tong University (Grant No. YG2014QN22).

## Data availability

The original data included in this study are presented in this article. Further inquiries can be directed to the corresponding author.

## Abbreviations

ANOVA: Analysis of Variance
BCA: Bicinchoninic Acid
CDX: Cell-Derived Xenograft
cDNA: Complementary DNA
CTLA-4: Cytotoxic T-Lymphocyte–Associated Protein 4
DEG(s): Differentially Expressed Gene(s)
DAPI: 4′,6-Diamidino-2-Phenylindole
EMT: Epithelial–Mesenchymal Transition
ESCC: Esophageal Squamous Cell Carcinoma
FBS: Fetal Bovine Serum
FITC: Fluorescein Isothiocyanate
GAPDH: Glyceraldehyde-3-Phosphate Dehydrogenase
GO: Gene Ontology
H&E: Hematoxylin and Eosin
HER2: Human Epidermal Growth Factor Receptor 2
HRP: Horseradish Peroxidase
HSP90: Heat Shock Protein 90
HSP90AA1: Heat Shock Protein 90 Alpha Family Class A Member 1
IHC: Immunohistochemistry
IF: Immunofluorescence
KEGG: Kyoto Encyclopedia of Genes and Genomes
mRNA: Messenger RNA
NK cells: Natural Killer cells
O-GlcNAc: O-linked N-acetylglucosamine
PBS: Phosphate-Buffered Saline
PCNA: Proliferating Cell Nuclear Antigen
PD-1: Programmed Cell Death Protein 1
PD-L1 (CD274): Programmed Death Ligand 1
PDX: Patient-Derived Xenograft
PI: Propidium Iodide
PRMT5: Protein Arginine Methyltransferase 5
PTM(s): Post-Translational Modification(s)
qPCR: Quantitative Polymerase Chain Reaction
RMA: Robust Multi-array Average
ROC: Receiver Operating Characteristic
RNAseq: RNA Sequencing
SD: Standard Deviation
SDMA: Symmetric Dimethylarginine
shRNA: Short Hairpin RNA
SPF: Specific Pathogen-Free
SDS-PAGE: Sodium Dodecyl Sulfate Polyacrylamide Gel Electrophoresis
SVR: Support Vector Regression
TCGA: The Cancer Genome Atlas
TPM: Transcripts Per Million
TRED: Transcriptional Regulatory Element Database
Tregs: Regulatory T cells
WB: Western Blot
WT: Wild Type

## Ethics Approval and Consent to Participate

This research was reviewed and approved by the Ethics Committee of Shanghai Chest Hospital (Approval No. KS(Y)-22005). All experimental procedures involving human subjects complied with institutional as well as national ethical regulations, following the principles outlined in the Declaration of Helsinki and its subsequent revisions or equivalent international standards. Prior to participation, written informed consent was obtained from all enrolled individuals and, when necessary, from their legal guardians.

## Consent for publication

Each participant provided written consent for the use of their anonymized data in publications derived from this study.

## Competing interest

The authors declare that they have no financial or personal conflicts of interest that could inappropriately influence, or be perceived to influence, the research described in this article.

## Notes

### Competing Interest Statement

The authors have declared no competing interest.

