## supplementary material for "PRMT5-Mediated Arginine Methylation of HSP90AA1 Drives Esophageal Squamous Cell Carcinoma Progression"

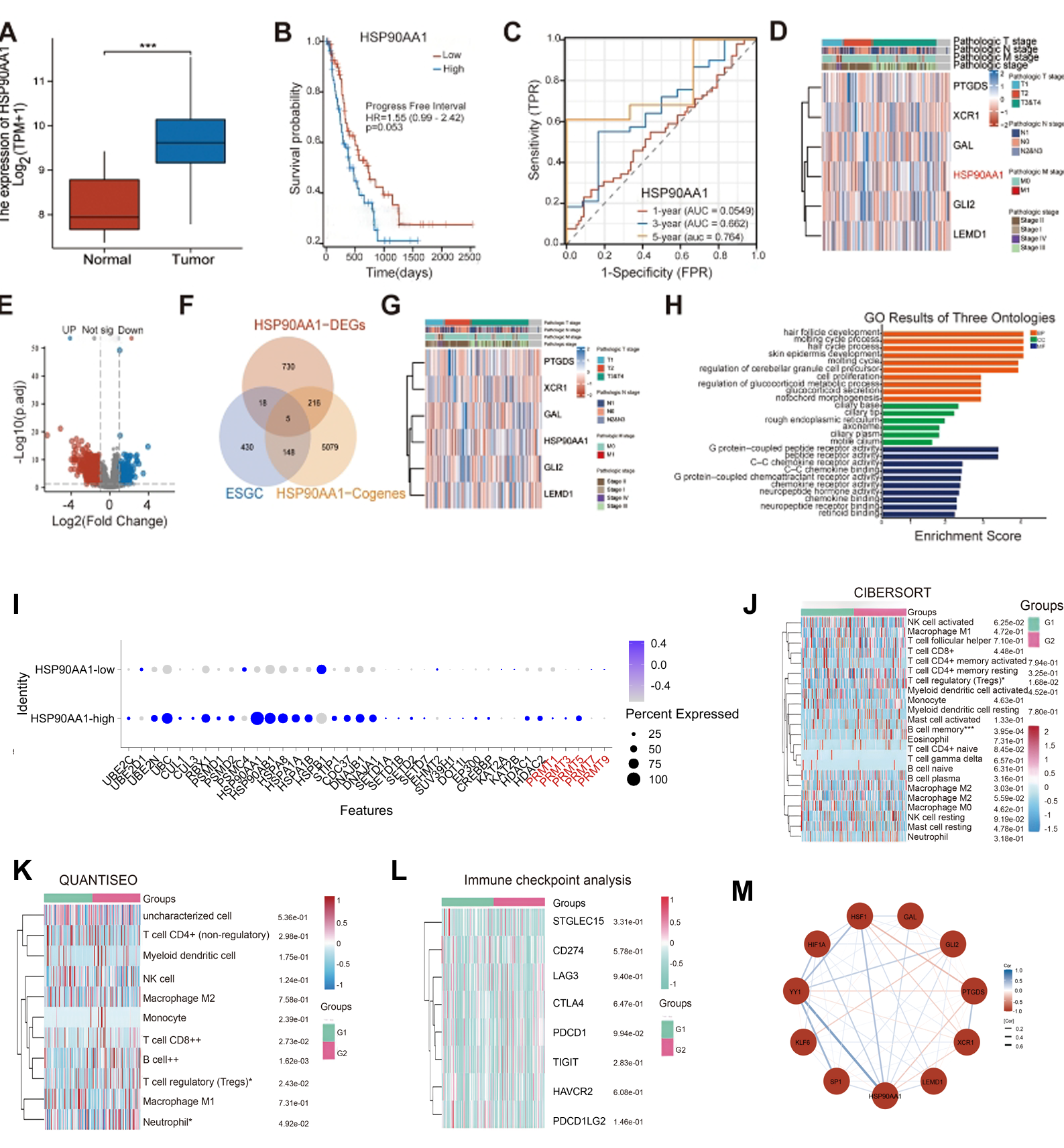

**Supplementary Figure 1. HSP90AA1-associated transcriptomic features, functional enrichment, immune contexture, and regulatory network in ESCC**

(A) Box plot showing HSP90AA1 expression in normal versus tumor samples.

(B) Kaplan–Meier analysis of progression-free survival for ESCC patients stratified by HSP90AA1 expression (high vs low).

(C) Time-dependent ROC curves evaluating the prognostic performance of HSP90AA1 (1-, 3-, and 5-year).

(D) Heatmap displaying expression patterns of the five intersecting genes (GAL, GLI2, PTGDS, XCR1, LEMD1) with clinicopathological annotations.

(E) Volcano plot of differentially expressed genes (DEGs) between HSP90AA1-high and HSP90AA1-low groups.

(F) Venn diagram showing the overlap among HSP90AA1-associated DEGs, ESCC-related genes, and HSP90AA1 co-expressed genes, identifying five shared genes (GAL, GLI2, PTGDS, XCR1, LEMD1).

(G) Heatmap illustrating the association of the five shared genes with clinicopathological features.

(H) GO enrichment results (BP/CC/MF) for the five shared genes.

(I) Dot plot showing expression of a curated post-translational modification (PTM)-related gene set in HSP90AA1-high versus HSP90AA1-low groups (dot size indicates percent expressed; color indicates average expression).

(J) Immune cell infiltration estimated by CIBERSORT, shown as a heatmap across samples/groups.

(K) Immune cell infiltration estimated by quanTIseq, shown as a heatmap across samples/groups.

(L) Heatmap of immune checkpoint gene expression across groups.

(M) Predicted transcriptional regulatory network for the five shared genes, visualized as a transcription factor–target interaction graph.

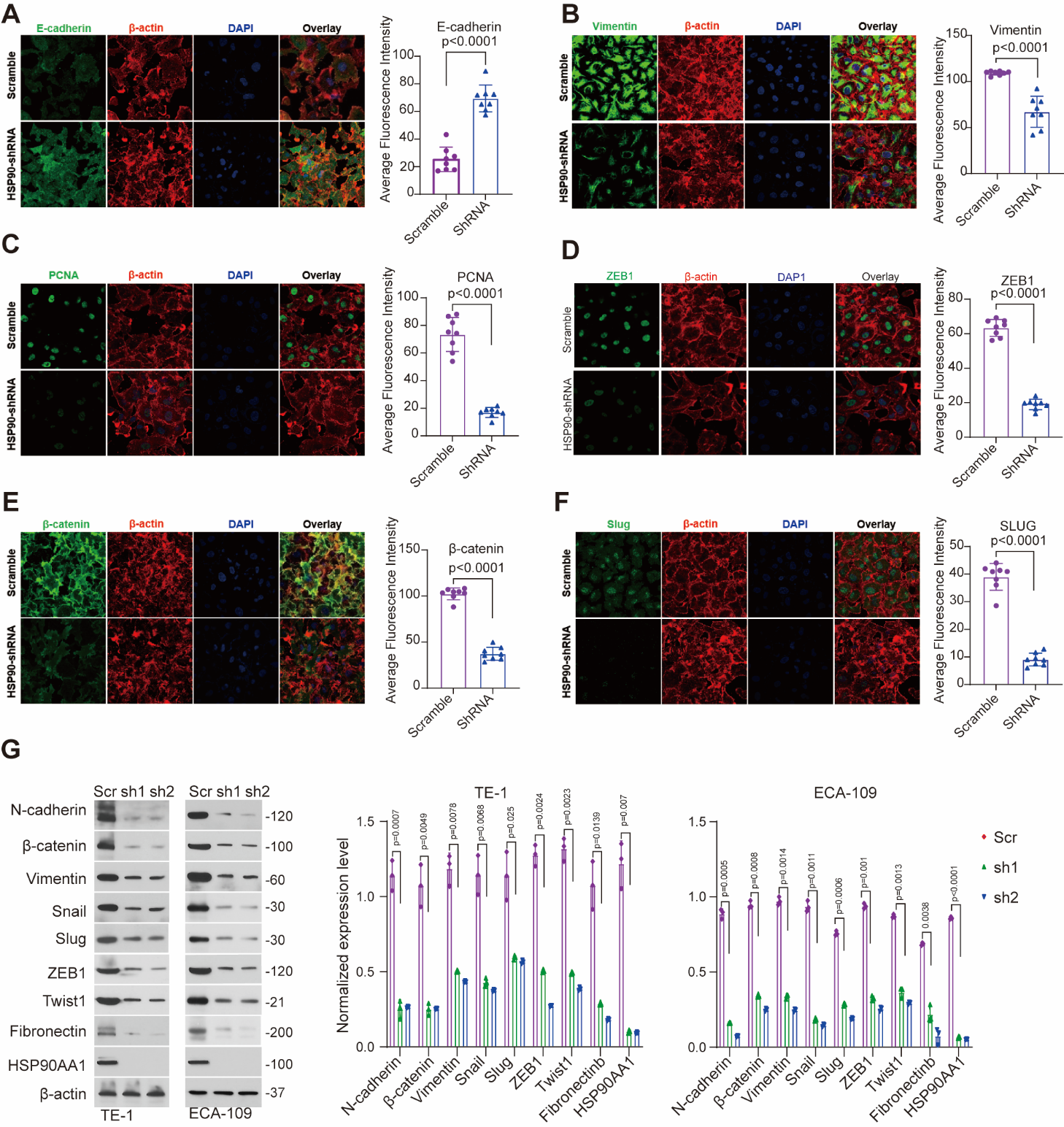

**Supplementary Figure 2. HSP90aa1 drives EMT in ESCC**

(A–F) Immunofluorescence (IF) staining in ESCC cells transfected with shRNA/HSP90AA1 or scramble control. Markers include E-cadherin (epithelial), Vimentin (mesenchymal), PCNA (proliferation), ZEB1, β-catenin, and Slug (EMT transcription factors).

(G) Western blot analysis of EMT and invasion/metastasis markers, including N-cadherin, Vimentin, Fibronectin, Snail, Slug, ZEB1, Twist1, and β-catenin, in ESCC cells following HSP90AA1 knockdown. β-actin was used as a loading control.

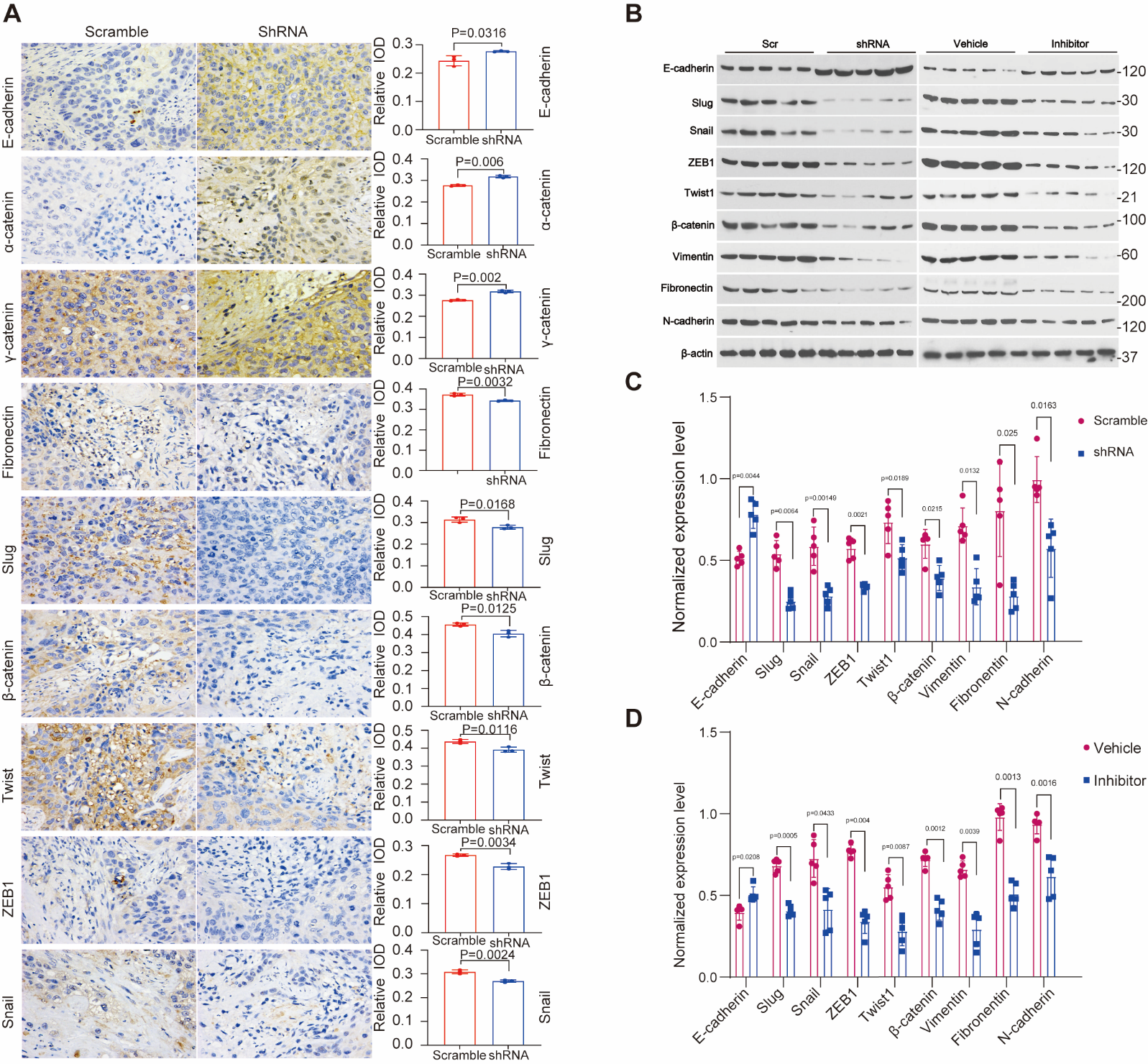

**Supplementary Figure 3. HSP90 inhibition or HSP90AA1 knockdown alters epithelial and mesenchymal marker expression**

(A) Immunohistochemical (IHC) staining showing increased epithelial markers (E-cadherin, α-catenin, γ-catenin) and decreased Fibronectin, ZEB1, Twist1, Snail, Slug, and β-catenin in ESCC cells treated with shRNA targeting HSP90AA1.

(B, C, D) Western blot analysis confirmed that HSP90 inhibition or HSP90AA1 knockdown upregulates epithelial markers while downregulating mesenchymal and pro-metastatic proteins.
